# Metastability in EEG phase synchronization networks is associated with autistic traits in a neurotypical cohort

**DOI:** 10.64898/2026.08.09.743722

**Authors:** Mebuki Izumiya, Yuka O Okazaki, Keiichi Kitajo

**Affiliations:** Department of Physiological Sciences, School of Life Sciences, The Graduate University for Advanced Studies, SOKENDAI, Okazaki, Aichi 444-8585 Japan; Division of Neural Dynamics, Department of System Neuroscience, National Institute for Physiological Sciences, National Institutes of Natural Sciences, Okazaki, Aichi 444-8585 Japan

## Abstract

Metastability is a fundamental dynamical property of large-scale brain networks and reflects the capacity of the brain to flexibly reorganize transient coordination patterns. In this study, we investigated whether metastable properties of resting-state electroencephalographic (EEG) phase synchronization networks are associated with individual differences in autistic traits. Resting-state EEG data from 88 neurotypical adults were analyzed using two complementary metrics: synchrony coalition entropy (SCE), which quantifies the diversity of transient phase synchronization patterns, and the metastability index (MSI), which quantifies temporal variance in global phase synchronization. SCE showed frequency-specific associations with the Autism-Spectrum Quotient (AQ) attention-switching subscore at 18–24 Hz and the communication subscore at 4–8 Hz, suggesting that frequency– and network-specific patterns of metastable synchronization are associated with distinct aspects of autistic traits. In contrast, MSI showed a modest association with the social-skill subscore in the lower-beta range, but this effect did not survive a cluster-based permutation test. This exploratory observation suggests that global synchronization variability may capture a weaker, complementary aspect of trait-related metastable dynamics. These findings suggest that, within a neurotypical population, individual differences in autistic traits may be more sensitively captured by the repertoire of transient phase synchronization patterns, as indexed by SCE, than by global phase synchronization variability, as indexed by MSI. Moreover, the associations of distinct AQ subscores with SCE in different frequency ranges suggest that different dimensions of autistic traits may be related to metastable network dynamics operating at different temporal scales.

**Author Summary:** The brain constantly coordinates activity across many regions, and this coordination changes over time rather than remaining constant. Understanding these dynamic patterns is important for explaining individual differences in cognition and behavior. In this study, we focused on a dynamical property called “metastability,” which describes how brain activity flexibly shifts between different patterns of coordination. Instead of remaining in a stable state, the brain repeatedly forms and dissolves coordinated activity across regions. We analyzed brain signals recorded with resting-state electroencephalography (EEG) and examined whether these dynamic patterns were related to individual differences in autistic traits. We found that different aspects of time-varying coordination were linked to different dimensions of autistic traits in a neurotypical population. These findings suggest that examining how brain activity changes over time, rather than relying only on time-averaged measures, can reveal neural features associated with individual differences in autistic traits. Our study highlights metastability as a useful concept for understanding the flexible and dynamic nature of human brain function.

## Introduction

Transient phase synchronization of neural activity is important for mediating the large-scale integration of distributed brain activity. Such transient phase synchronization networks are thought to support the dynamical integration of distributed information processing that underlies cognitive processes [1]. Importantly, these networks do not remain fixed in a single configuration but instead exhibit flexible transitions among multiple coordination states. This transient and adaptive regime is essential for efficient and adaptive cognitive function [2]. In this study, we focused on metastable phase synchronization networks and investigated how their dynamical properties, measured using resting-state electroencephalography (EEG), relate to individual psychological traits.

Metastability refers to a regime of dynamical systems in which multiple coordination states coexist as weakly attracting states. In such a regime, the system temporarily dwells near each state without converging to a single stable equilibrium, and spontaneously transitions among them over time [3]. Importantly, metastability does not denote the transition events themselves but rather is a property of the system’s dynamical landscape, characterized by the absence of a globally stable attractor and the presence of transiently stable coordination patterns [4]. In contrast to multistability, in which transitions between stable attractors are typically driven by external perturbations or noise, metastable systems intrinsically exhibit ongoing switching because of the balance between coupling and autonomy among system components. This balance allows the components to synchronize transiently while retaining their capacity for independent dynamics, thereby enabling both integration and segregation within the brain system [2, 3]. Thus, metastability captures a regime between full synchronization and complete independence. In the context of brain dynamics, metastability has been proposed as a fundamental organizing principle supporting flexible information processing and cognitive adaptability. Neural systems operating in a metastable regime can rapidly reconfigure their functional networks, allowing the brain to explore a rich repertoire of coordination patterns without being locked into rigid states [2, 4–6].

Phase synchronization is thought to support the integration and processing of information across distributed brain regions [1]. Consequently, the primary objective of our study was to apply and extend existing metastability-related metrics to spatially resolved EEG phase synchronization networks and individual psychological traits, with a specific focus on autistic traits that are continuously distributed in the general population and can be assessed using the Autism-Spectrum Quotient (AQ). Autism spectrum disorder (ASD) is a developmental disorder characterized by difficulties in social interaction and communication and restricted or repetitive behaviors [7]. Atypical sensory responsiveness, including hyperresponsiveness, hyporesponsiveness, and sensory seeking, is also frequently observed in ASD [7]. Although ASD is a categorical diagnosis, autistic characteristics are widely considered to lie on a continuum across individuals and can be observed in both clinical and neurotypical populations [8–10].

Further motivation for this study comes from the methodological gap in quantifying the metastable dynamics of large-scale EEG phase synchronization in a spatially resolved manner. Previous studies have demonstrated that dynamical properties derived from EEG—computed from scalp topographies, amplitude-based activity, or phase-based coupling—can capture meaningful variations across brain states and individuals [11–13]. EEG studies have shown that the temporal structure of phase synchrony is altered in patients with ASD [14]. However, studies that jointly characterize the global and spatially specific metastability of phase synchronization networks and relate them to autistic traits remain limited.

In this study, we quantified resting-state metastable dynamics using two complementary metastability metrics derived from phase synchronization networks: the metastability index (MSI), which reflects the temporal variability of global synchronization, and synchrony coalition entropy (SCE), an entropy-based index of the diversity of transient synchronization coalitions originally introduced in an information-theoretic framework (e.g., [15]) and applied to EEG in a prior study [16]. Importantly, we examined the associations between MSI and SCE values and AQ scores and used topographic maps to characterize the spatial distribution of channel-wise SCE. By focusing on how metastable phase synchronization dynamics are related to autistic traits, we aimed to clarify trait-relevant signatures of resting-state network dynamics, providing a step toward a biologically grounded, individualized characterization of neurocognitive diversity.

## Materials and Methods

### Data acquisition

#### EEG dataset

We analyzed our previously published resting-state EEG dataset available on OSF [13] (https://doi.org/10.17605/OSF.IO/29QB5). EEG data were obtained from 132 participants (66 females; mean age ± SD, 24.0 ± 5.0 years). All participants provided informed consent before starting the experiment. The study was approved by the RIKEN Ethics Committee (Wako3 26–24).

For the EEG recordings, an EEG amplifier (BrainAmp MR Plus, Brain Products GmbH, Gilching, Germany) with a 63-channel EEG cap (EasyCap, EASYCAP GmbH, Herrsching, Germany) was used. The sampling rate was 1000 Hz, and the low and high cutoff frequencies of the online filter were 0.016 and 250 Hz, respectively. Electrodes were placed over the scalp according to the 10–10 system. AFz was set as the ground electrode, the reference electrode was placed on the left earlobe, and offline re-referencing was conducted using the average potentials of the left and right earlobes. All recorded scalp electrodes were included in the analysis. Participants were asked to rest and close their eyes during the 3-min EEG recording.

Of the 132 participants in the original dataset, 88 had resting-state EEG recordings and AQ questionnaire data available for analysis. Therefore, the main cross-sectional analysis included 88 participants. None of the participants were selected or excluded based on their AQ scores. EEG data quality control followed the procedures described in the original dataset publication. Their ages ranged from 20 to 47 years (45 females; mean age ± SD, 24.4 ± 5.6 years). The test–retest reliability analysis included a subsample of 32 participants (13 females; mean age ± SD, 25.2 ± 6.5 years) who completed a follow-up EEG recording after a mean interval of 101 days.

#### Autism-Spectrum Quotient (AQ)

Autistic traits were assessed in 88 participants using the Japanese version of the AQ [17–19]. Detailed descriptive statistics and AQ score distributions for the full sample and the test–retest subsample are provided in S1 Table and S1 Fig. The total AQ scores ranged from 5 to 41 (mean ± SD, 19.28 ± 7.40; median, 19). The distribution of total AQ scores did not significantly deviate from normality (*p* = 0.206, Shapiro–Wilk test), with the largest proportion of participants falling in the 20–25 score range. The AQ subscores were as follows: *social skill*, mean ± SD, 4.19 ± 2.68; *attention switching*, 4.24 ± 2.02; *attention to detail*, 4.25 ± 2.18; *communication*, 2.98 ± 2.33; and *imagination*, 3.63 ± 2.09.

In the test–retest subsample (*n* = 32), the total AQ scores ranged from 6 to 33 (mean ± SD, 20.31 ± 7.70; median, 21.5). The distribution of AQ scores did not significantly deviate from normality (*p* = 0.155, Shapiro–Wilk test). The AQ subscores were as follows: *social skill*, mean ± SD, 4.81 ± 2.84; *attention switching*, 4.41 ± 2.00; *attention to detail*, 3.88 ± 2.43; *communication*, 2.84 ± 2.15; and *imagination*, 4.38 ± 2.22.

### Analysis

All analyses were conducted using custom-written MATLAB scripts (Version R2024a, MathWorks Inc.), EEGLAB (Version 2022.1) [20], and the FieldTrip toolbox [21]. Custom code for calculating MSI and SCE is described in the Code availability section. The analysis pipeline is shown in Fig. 1.

**Fig 1.**
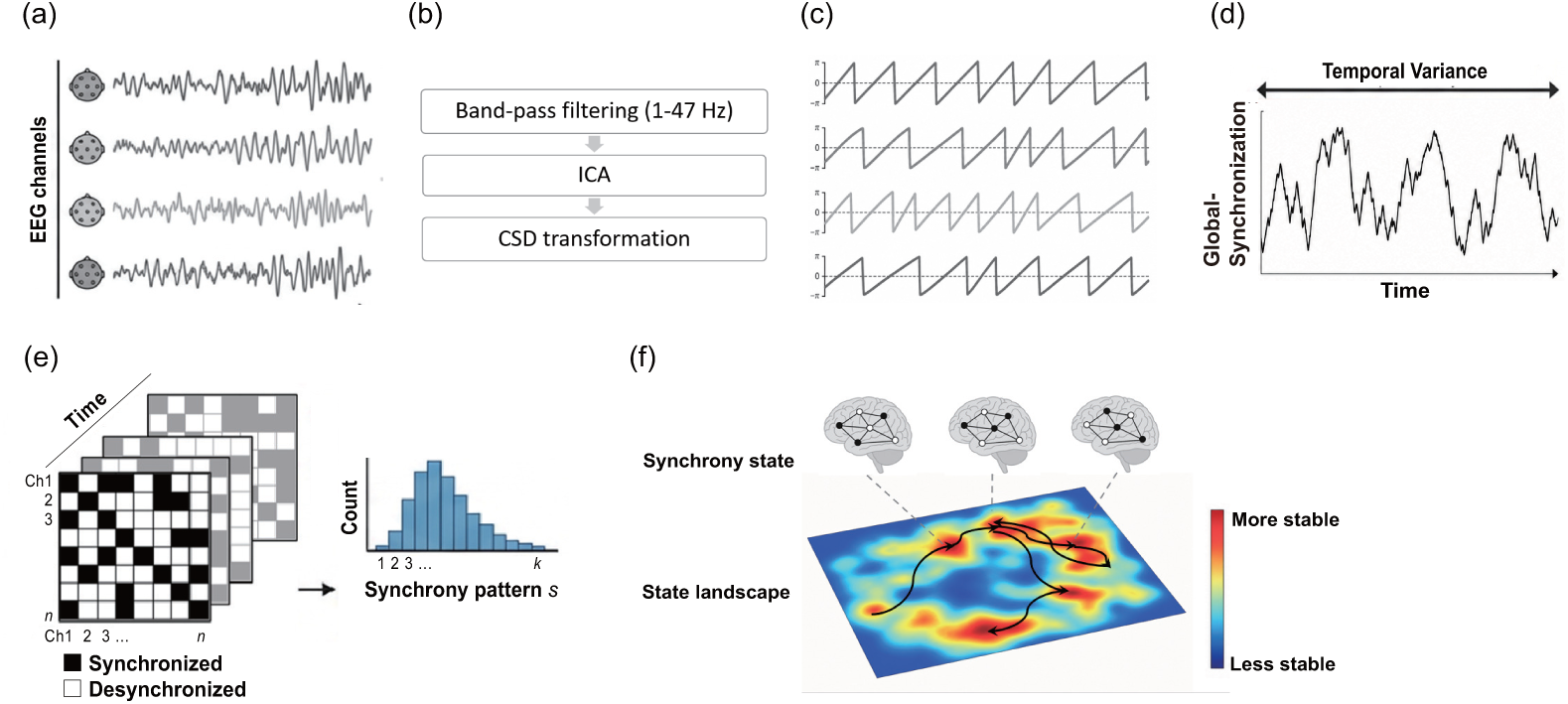
Analysis overview of this study. (a) Raw EEG signals recorded from multiple channels. (b) EEG preprocessing including bandpass filtering (1-47 Hz), independent component analysis (ICA), and current source density (CSD) transformation, which are described in the next section. (c) Instantaneous phase time series extracted from each EEG signal using wavelet-based phase decomposition. The phase evolves continuously over time and wraps between *π* and *π*. (d) Metastability Index (MSI), quantified as the temporal variance of the global synchronization level, reflecting fluctuations in large-scale phase coordination over time. Larger temporal fluctuations correspond to higher MSI values. (e) Synchrony coalition entropy (SCE), quantifying the diversity of binary phase synchronization patterns across channel pairs. (f) Conceptual schematic illustrating how metastable state-space dynamics involve continuous transitions among multiple network states over an underlying state landscape. This dynamical regime enables both integration and segregation of neural activity across time. This is a conceptual schematic and does not represent an empirically estimated energy landscape.

**Fig 2.**
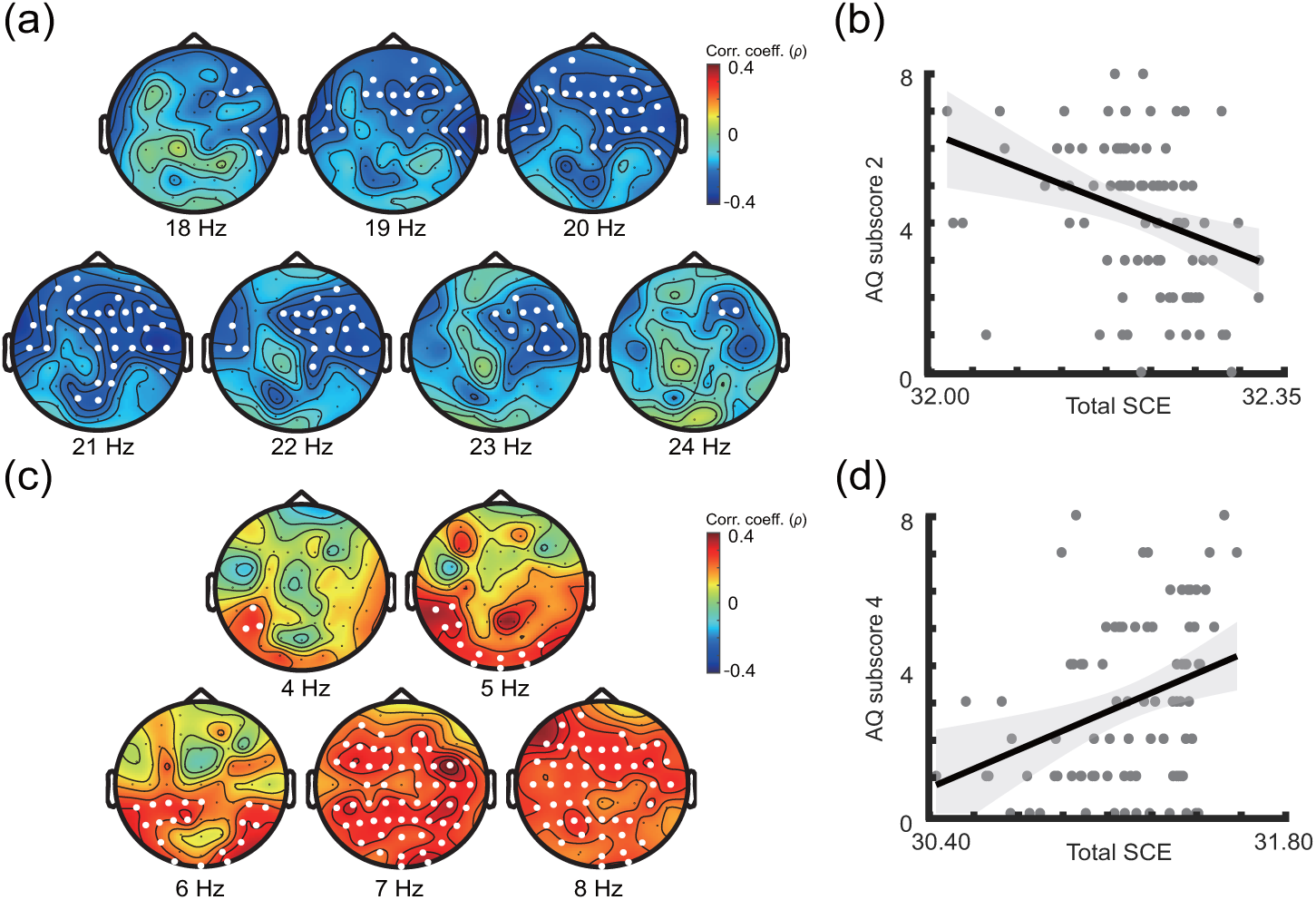
Frequency-specific associations between SCE and AQ subscores. Topographic maps show Spearman’s rank correlation coefficients between SCE and AQ (a) *attention switching* and (c) *communication* scores across 1–47 Hz. Electrodes belonging to significant clusters are marked with white dots. Scatter plots show the relationship between SCE summed within each significant cluster and the corresponding AQ (b) *attention switching* and (d) *communication* scores. Each point represents an individual participant, and solid lines indicate linear fits for visualization purposes. Statistical significance was assessed using a two-tailed cluster-based permutation test with 5,000 permutations and a cluster-forming threshold of *p <* 0.05.

#### EEG preprocessing

EEG data were bandpass filtered from 1 to 47 Hz using the EEGLAB legacy function eegfilt. The function implements a two-pass zero-phase finite impulse response (FIR) filter designed using a least-squares method. The lower and upper passband edges were set to 1 and 47 Hz, respectively. With a sampling rate of 1,000 Hz and a lower cutoff frequency of 1 Hz, the default filter order corresponded to approximately 3,000 points. Filtering was applied to the continuous data before subsequent time-frequency decomposition.

Next, independent component analysis (ICA) was performed in EEGLAB [20, 22], and ICLabel [23] was used to classify independent components. ICA was performed using the EEGLAB function pop_runica with the default Infomax ICA algorithm (runica). Artifactual independent components classified as eye or muscle artifacts with a probability greater than 0.8 were removed.

Current source density (CSD) transformation was applied using the CSD toolbox [24, 25] to implement the spherical spline surface Laplacian. The electrode positions were defined according to the 10–10 system using the 3D electrode coordinates provided by the EasyCap montage, and the default smoothing parameter was used. This transformation was performed before the phase extraction to reduce the influence of volume conduction and reference-dependent effects.

Time–frequency decomposition was performed using a complex Morlet wavelet [26, 27] to extract the instantaneous phase of the signals. The analytic wavelet *ψ_f_* (*t*) was defined as a Gaussian-windowed complex exponential:

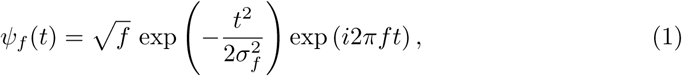

where *f* denotes the central frequency and *σ_f_* determines the temporal width of the Gaussian envelope. The temporal width was defined as

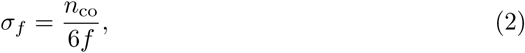

where *n*_co_ denotes the number of wavelet cycles. Because *σ_f_* is inversely proportional to *f*, the scaling factor √*f* makes the wavelet energy independent of frequency up to a frequency-independent multiplicative constant. The complex-valued wavelet-transformed signal was obtained by convolution of the EEG signal *s*(*t*) with the wavelet:

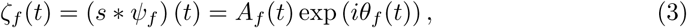

where *A_f_* (*t*) and *θ_f_* (*t*) represent the instantaneous amplitude and phase at frequency *f*, respectively. Because the present study focused on phase-based measures of connectivity and metastability, only the instantaneous phase *θ_f_* (*t*) was used in subsequent analyses.

#### Metastability index (MSI)

Shanahan [15] proposed metastability metrics using a community-structured coupled oscillator model based on the Kuramoto model [28], a standard mathematical framework for synchronization phenomena. In Shanahan’s study, metastability was maximized in a given parameter region using a coupled oscillator model. In this study, we used two previously proposed metastability-related metrics. The metastability index (MSI) was calculated as follows. The Kuramoto order parameter, *r*(*t*), which quantifies the degree of phase synchronization, is given by:

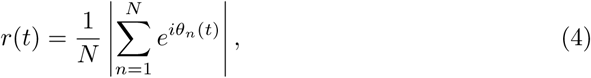

where *N* is the number of oscillators, corresponding to the number of EEG channels, and *r*(*t*) represents the level of instantaneous synchronization. The oscillators are fully synchronous when *r*(*t*) is 1 and completely asynchronous when *r*(*t*) is 0. The temporal variance of the order parameter *r*(*t*) was defined as the MSI in the current study. MSI reflects the temporal variability in the global synchronization level. A high MSI indicates greater temporal variability in large-scale phase coordination across EEG channels. Thus, this variance estimate is a measure of the metastability of the entire system, as follows:

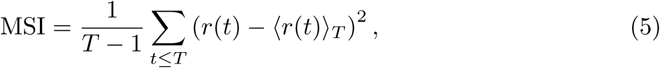

where *T* is the total number of time points over the recording period (180,000 points). MSI is the temporal variance of the order parameter *r*(*t*).

#### Synchrony coalition entropy (SCE)

Prior research has proposed that the extent of metastability can be evaluated through the computation of synchrony coalition entropy (SCE) [15, 16]. SCE was calculated from the temporal patterns of pairwise phase differences. The procedure for computing channel-wise SCE is illustrated in Fig 1. Let *M* denote the total number of electrode channels. The phase difference between channel *i* and channel *j* was computed as follows:

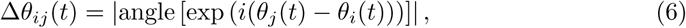

For each reference channel *i*, the phase differences with remaining *M* − 1 channels were binarized as follows:

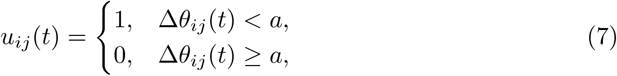

where *a* is the phase difference threshold. Let **u_i_**(*t*) 0, 1 *^M−^*^1^ denote the binary synchrony vector, or coalition pattern, for reference channel *i* at time *t*. For each reference channel *i*, the binary vector **u_i_**(*t*) was constructed from the *M* − 1 pairwise phase relationships. The probability *p*(**s***_k_*) was then estimated as the relative frequency of occurrence of the synchrony coalition pattern **s***_k_*, where *k* indicates the distinct patterns observed over time. SCE was computed using the Shannon entropy of this distribution. Thus, the channel-wise SCE is denoted as

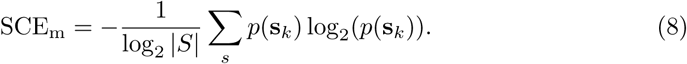

Here, *S* denotes the number of possible distinct coalition patterns. For *M* EEG channels, there are *S* = 2*^M−^*^1^ possible coalitions for each reference channel. SCE_m_ was retained as a channel-wise measure. For whole-brain summary analyses, oSCE was computed by averaging SCE_m_ across channels. For analyses requiring a single whole-brain summary value, we computed the average SCE across all channels, denoted as:

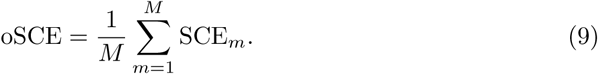

#### Statistical analysis

All statistical analyses were conducted across participants. Associations between EEG-derived metastability metrics and AQ scores were assessed using Spearman’s rank correlation coefficient, denoted as *ρ*. Spearman’s correlation was selected because it evaluates monotonic associations without assuming linear relationships or normal variable distributions. Correlations were computed between each EEG-derived measure and the total AQ score or each subscore.

Test–retest reliability was assessed using a one-way random-effects intraclass correlation coefficient, ICC(1, 1), for each frequency bin and metastability measure in participants who underwent two recording sessions (subsample *n* = 32). The model was written as

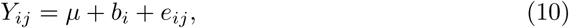

where *Y_ij_*is the value for participant *i* at session *j*, *µ* is the grand mean, *b_i_*is the participant-specific random effect, and *e_ij_* is the residual error. The variance of the participant-specific random effect was denoted as *τ* ^2^, and the residual variance was denoted as *τ* ^2^. Population-level ICC(1, 1) was defined as the proportion of the total variance attributable to between-participant differences.

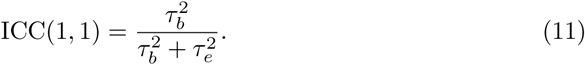

For frequency-wise reliability analyses, p-values were corrected for multiple comparisons across frequencies using the false discovery rate (FDR) procedure.

Cluster-based permutation tests were performed using the FieldTrip toolbox to assess the sensor-frequency associations between the SCE and AQ scores. SCE values were computed for each participant, electrode, and frequency bin and arranged in a FieldTrip-compatible format. For each electrode-frequency point, the association between SCE and AQ scores was evaluated across participants using Spearman’s rank correlation. In the FieldTrip framework, this was implemented by rank-transforming the variables across participants before applying correlation statistics. The resulting Spearman’s rank correlation coefficients, *ρ*, were retained as effect-size estimates.

Statistical significance was assessed using a two-tailed Monte Carlo cluster-based permutation test with 5,000 random permutations. Clusters were formed across electrodes and frequency bins using a cluster-forming threshold of *p <* 0.05. Spatial adjacency was defined based on the inter-electrode proximity, with a minimum of two neighboring electrodes required for cluster formation. Cluster-level significance was evaluated using the *maxsum* cluster statistic, and family-wise error correction was applied at the cluster level with a significance threshold of *p <* 0.05. Positive and negative clusters were evaluated separately.

The design matrix contained a single subject-level regressor: the AQ score. Correlation analysis was performed independently for each electrode and frequency bin before cluster-level correction. Clusters that survived the cluster-level correction were visualized as topographic maps over the corresponding frequency ranges. These topographic maps were used to visualize the spatial distribution of significant sensor-frequency clusters, and statistical inferences were based on the cluster-level permutation results.

For MSI, frequency-wise associations with AQ scores were examined using Spearman’s rank correlation. As the MSI–AQ associations did not survive cluster-level correction, these results were treated as exploratory. Additionally, electrodes-of-interest analyses based on SCE-defined clusters were performed as post-hoc descriptive analyses and were not used for confirmatory statistical inference.

The phase difference threshold for SCE binarization was determined using the test-retest reliability subsample (*n* = 32), independent of the main AQ association analysis. Candidate thresholds were evaluated based on the test–retest reliability of the SCE, and *a* = 1.2 rad was selected as the primary threshold. The threshold-dependent reliability profiles are shown in S2 Fig.

#### Uniform Manifold Approximation and Projection (UMAP) for visualization of individual metastability

We visualized individual metastable dynamics using the binarized phase-difference matrices (0/1) obtained from the SCE analysis. Prior to nonlinear dimensionality reduction, principal component analysis (PCA) was applied to the original high-dimensional binary phase-difference matrices to reduce noise, redundancy, and computational complexity. The number of retained principal components was determined such that the cumulative explained variance exceeded 80%, thereby preserving most of the variance in the original data while reducing the dimensionality of the feature space.

Subsequently, UMAP was used to embed the PCA-reduced data into a two-dimensional space for visualization [29]. UMAP is a nonlinear dimensionality reduction method designed to preserve local neighborhood relationships while approximating the global structure of the underlying data manifold. In this study, UMAP was implemented in MATLAB using the publicly available UMAP package [30]. A cosine distance metric was used to quantify similarity between observations in the PCA-reduced feature space. The number of nearest neighbors was set to 10 to emphasize the local neighborhood structure, and the minimum distance parameter was set to 0.01 to allow similar states to be embedded closely in the low-dimensional space. Fixed random seeds were used to ensure reproducibility.

This dimensionality reduction procedure was used for exploratory visualization of complex EEG-derived network dynamics. Because UMAP analysis was applied to data from a limited number of participants, the resulting embeddings were interpreted descriptively and were not used for statistical inference.

## Results

### The relationship between SCE and AQ

We examined sensor–frequency clusters of Spearman’s rank correlations between SCE and AQ scores, including the total AQ score and each AQ subscore. Significant associations with SCE were identified for the *attention switching* and *communication* subscores.

For *attention switching*, a significant negative cluster was identified in the upper-beta range (18–24 Hz; cluster statistic = −324.4; cluster-level *p* = 0.032), indicating that higher SCE values were associated with lower AQ *attention switching* scores. For *communication*, a significant positive cluster was observed in the theta range (4–8 Hz; cluster statistic = 356.0; cluster-level *p* = 0.036), indicating that higher SCE values were associated with higher AQ *communication* scores. The full frequency-wise correlation profiles between AQ scores and whole-brain summary metrics are shown in S3 Fig.

### The relationship between MSI and AQ

We examined the frequency-specific associations between AQ scores and MSI using Spearman’s rank correlation across frequencies.

The correlation spectrum exhibited a local peak around 17–19 Hz, with consistently positive correlations at these frequencies (17 Hz: *ρ* = 0.215, *p* = 0.0438; 18 Hz: *ρ* = 0.224, *p* = 0.0355; 19 Hz: *ρ* = 0.220, *p* = 0.0395; uncorrected, Fig 3(a)). However, cluster-based permutation testing across frequencies did not reveal any significant clusters.

**Fig 3.**
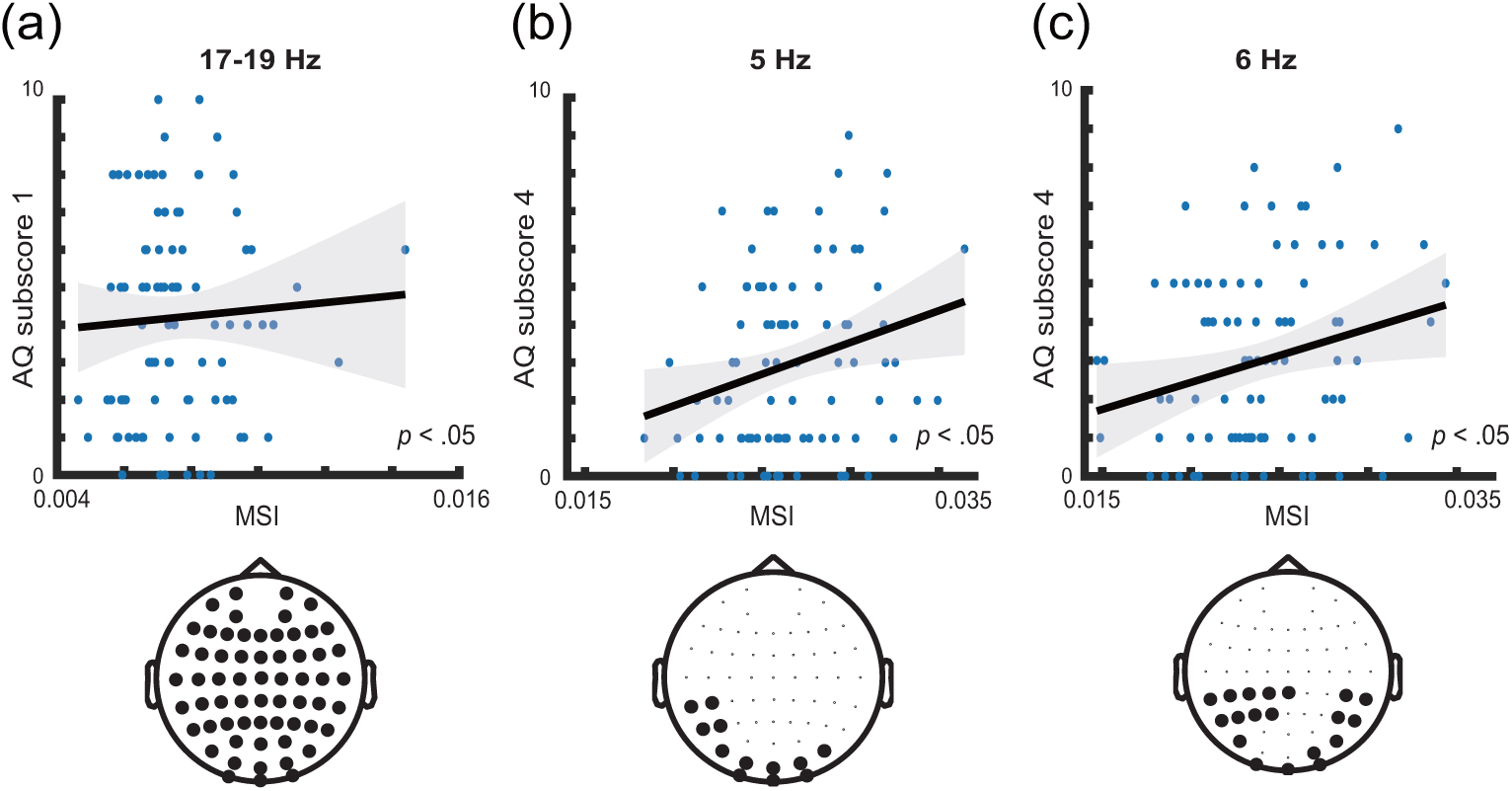
Frequency-specific associations between MSI and AQ subscores. (a) Correlation between full-channel MSI averaged across the 17–19 Hz range and the AQ *social skill* subscore. Correlations between EOI-restricted MSI and the AQ *communication* subscore at (b) 5 Hz and (c) 6 Hz. EOI-restricted MSI was calculated using only electrodes that showed significant channel-wise SCE–AQ correlations within the corresponding SCE-defined frequency cluster. Bottom topographies indicate the EOI electrode sets used for MSI recalculation in each frequency cluster. Solid lines represent linear fits with 95% confidence intervals shown in gray.

To further examine whether channels contributing to significant SCE–AQ associations also accounted for MSI–AQ relationships, we performed an exploratory electrodes-of-interest (EOI)-restricted MSI analysis. For each frequency cluster showing significant SCE–AQ correlations, the electrodes exhibiting significant channel-wise SCE–AQ correlations within that cluster were defined as EOIs. The MSI was then recalculated using only the phase signals from these EOIs, and Spearman’s rank correlation coefficients were computed between the EOI-restricted MSI and the corresponding AQ subscore.

However, the EOI-restricted MSI did not consistently reproduce the MSI–AQ associations observed in the full-channel analysis. While some frequency clusters retained nominal positive correlations, they did not reach statistical significance despite being derived from electrodes that showed robust SCE–AQ associations. For example, EOI-restricted MSI showed nominal positive correlations with AQ *communication* at 5 Hz (*ρ* = 0.225, *p* = 0.035) and 6 Hz (*ρ* = 0.213, *p* = 0.046), but these effects should be interpreted descriptively because the EOIs were defined post hoc based on the SCE–AQ results.

These results suggest that channels informative for SCE do not necessarily preserve the AQ-related variability captured by the MSI, indicating that the two metrics may partially reflect distinct aspects of metastable EEG network dynamics.

### Test-retest reliability for metastability index (MSI) and synchrony coalition entropy (SCE)

To assess test-retest reliability, we computed the ICC(1, 1) values across frequencies (1–47 Hz) using follow-up EEG data from 32 participants collected after a mean interval of 101 days. As shown in Fig 4, the MSI showed relatively high reliability, mainly in the delta to lower beta range (1–20 Hz), whereas SCE showed high reliability across a broader frequency range, extending from 2 to 47 Hz. Several frequency bins remained significant after FDR correction. Overall, the test–retest reliability analyses suggested that both MSI and SCE can be quantified reproducibly across sessions in resting-state EEG, indicating that these metrics capture the relatively stable properties of metastable phase synchronization networks.

**Fig 4.**
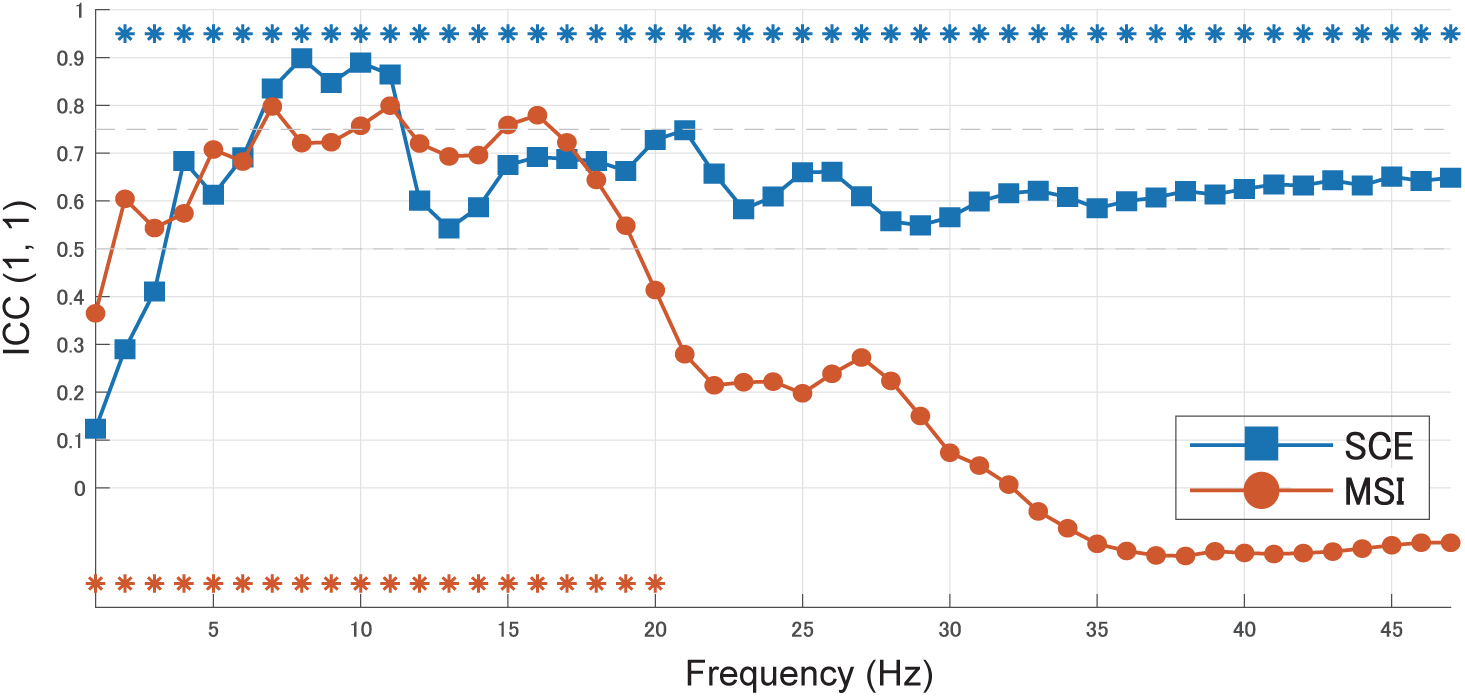
Test-retest reliability of MSI (orange) and SCE (blue) across frequencies (1–47 Hz) between the first and second recording sessions (*N* = 32). The plot shows the ICC(1, 1) values at each frequency. Asterisks indicate frequencies that remained significant after false discovery rate (FDR) correction (\**p <* 0.05).

### Individual representation of metastability

To visualize individual differences in the structure of high-dimensional EEG-derived network dynamics, we projected instantaneous network configurations onto a two-dimensional space using UMAP. For illustrative purposes, we selected the participants with the lowest and highest total AQ scores. In this embedding, nearby points represent similar network configurations in the original high-dimensional space, allowing recurrent states and transitions to be visualized.

The lowest-AQ participant exhibited a broad and complex distribution in the UMAP space, characterized by six distinct high-density regions in the density landscape (Fig 5a). Each high-density region corresponded to a recurrent metastable network configuration, indicating repeated visits to multiple distinct network states over time. The spatial separation among the high-density regions and the trajectories connecting them (Fig 5c) suggest a rich exploration of the underlying low-dimensional state space, reflecting frequent transitions among metastable configurations. In contrast, the highest-AQ participant showed a compact distribution dominated by a single cluster with one prominent peak and another weak peak in the UMAP space (Fig 5b). The trajectories of the network states remained confined to a limited region (Fig 5d), indicating that the network dynamics were largely restricted to a single dominant metastable configuration. Transitions to alternative configurations were relatively infrequent in this illustrative example.

**Fig 5.**
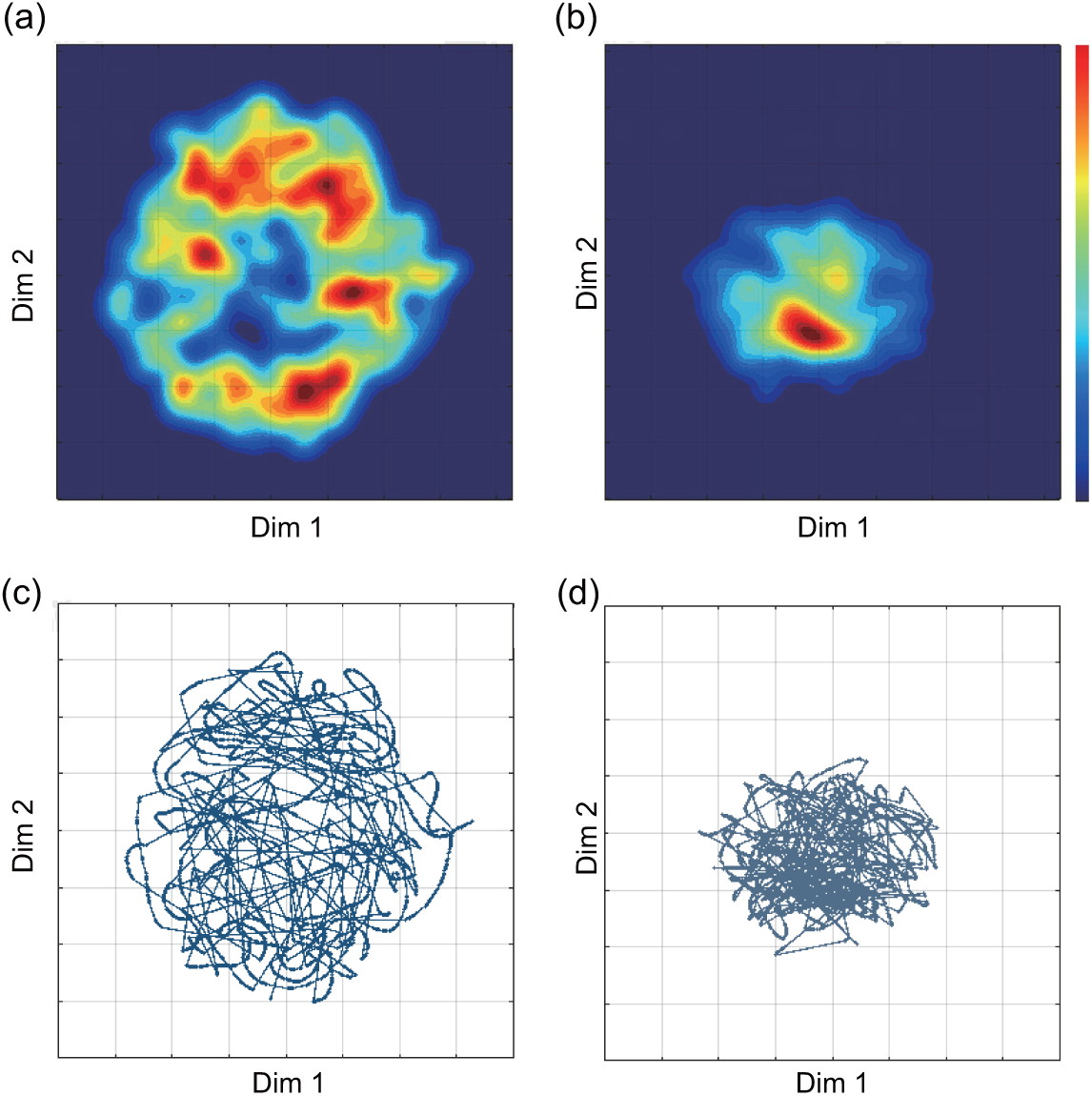
UMAP embeddings of phase synchronization dynamics in the lowest-AQ (left, total score was 5) and highest-AQ (right, total score was 37) individual participants. (a, b) The density map indicates the recurrence frequency of visited states, computed via kernel density estimation. (c, d) Lines indicate trajectories of the phase synchronization states across time. Notably, (a, c) the lowest-AQ individual showed multiple state clusters, whereas (b, d) the highest-AQ individual showed a more spatially restricted distribution.

## Discussion

In this study, we quantified EEG metastable phase synchronization networks using two metastability metrics: SCE, which captures the diversity of spatiotemporal phase synchronization patterns, and MSI, which reflects temporal fluctuations in global phase synchronization. The observed associations between these metrics and the AQ subscores provide insights into how dynamical brain network properties relate to individual differences in autistic traits within a neurotypical cohort.

First, we found that SCE showed cluster-based, frequency-specific associations with AQ subscores, highlighting the importance of spatiotemporal diversity in large-scale brain network dynamics. Unlike global synchrony metrics, SCE captures the repertoire of distinct phase synchronization patterns that emerge over time and therefore reflects the richness of metastable state configurations explored by the brain.

In the beta band (18–24 Hz), the negative correlation between SCE and AQ *attention switching* subscore suggests that reduced diversity of network states is associated with greater self-reported difficulty in shifting attention. This subscore can be interpreted as one aspect of the broader rigidity–flexibility dimension often discussed in autism, including difficulties in set shifting, perseveration, and restricted or repetitive patterns of behavior [17, 31–33]. From a dynamical systems perspective, a lower SCE value may indicate a more restricted accessible state space, in which the system tends to remain within a limited set of network configurations. Beta-band oscillations are known to stabilize ongoing cognitive states and support top-down control [34]; thus, a reduced SCE in this range may be consistent with a relatively stable regime in which transitions between metastable states are less frequent. This interpretation is broadly consistent with previous findings of perseveration and impaired set shifting associated with higher autistic traits, as well as theoretical accounts linking cognitive inflexibility to frontoparietal control networks [35, 36]. However, because the relationship between self-reported rigidity and laboratory metrics of cognitive flexibility is not always straightforward [31], the present findings should be interpreted as linking reduced metastable network diversity to attention-switching-related autistic traits rather than to cognitive flexibility as a whole.

In contrast, the positive correlation between the theta-band (4–8 Hz) SCE and the AQ *communication* subscore suggests that a greater diversity of network configurations at lower frequencies is associated with stronger self-reported communication-related autistic traits. Theta oscillations are implicated in long-range coordination and are thought to facilitate communication between distant cortical areas involved in language processing [37]. A higher SCE in the theta band may indicate a broader repertoire of metastable configurations, which may reflect altered variability in coupling and decoupling among communication-related networks. This interpretation aligns with previous findings that disruptions in theta-band coordination are associated with impaired communication in ASD [38], although the present results should be interpreted as a trait-related association in a neurotypical cohort rather than as evidence of ASD pathology.

Taken together, these findings suggest that SCE may index an aspect of brain dynamics related to the balance between stability and flexibility in network coordination. A higher SCE may correspond to a richer metastable landscape with multiple accessible states, whereas a lower SCE value may indicate a constrained dynamical regime with limited transitions. Importantly, the frequency-specific nature of these effects implies that different timescales of neural coordination contribute differently to cognitive functions, with beta-band dynamics relating more to cognitive stability and theta-band dynamics to integrative communication processes. Importantly, these findings should not be interpreted as biomarkers of clinical ASD because the present sample consisted of neurotypical adults, and autistic traits were assessed dimensionally using the AQ.

Associations involving MSI were comparatively weaker and more limited. We observed a modest relationship between MSI in the lower-beta band (17–19 Hz) and the *social skill* subscore. While beta-band activity has been linked to social cognition and large-scale network integration [39, 40], the present findings should be interpreted cautiously. MSI reflects global synchronization variability and may be less sensitive to spatially heterogeneous network configurations than SCE. Therefore, MSI may provide complementary but less specific information regarding the relationship between metastability and behavioral traits.

Notably, the frequency bands identified in this study differed from those reported in previous studies [13]. This discrepancy likely arises from methodological differences. While previous studies have employed PAC-based metastability metrics that incorporated both phase and amplitude dynamics, the current study focused on phase synchronization alone. These approaches capture distinct aspects of neural dynamics; PAC reflects cross-frequency interactions, whereas phase synchronization emphasizes transient large-scale coordination. Despite these differences, both frameworks highlight the importance of metastable dynamics in supporting flexible cognition.

To further illustrate the individual differences in metastable dynamics, we visualized the low-dimensional structure of the EEG state space using UMAP. The illustrative examples showed marked variability in the organization of metastable configurations across individuals; the lowest-AQ participant exhibited a distributed landscape with multiple distinct states, whereas the highest-AQ participant showed a more constrained structure dominated by a single configuration. Because this analysis was based on representative individuals rather than group-level statistical testing, these observations should be regarded as exploratory and hypothesis-generating. Nevertheless, they illustrate how individual differences in the repertoire of phase synchronization states may be visualized and may provide a useful framework for future studies of metastable brain dynamics.

It is important to note that MSI and SCE should be interpreted as empirical signatures of metastable dynamics rather than as direct measurements of metastability. Metastability is formally a property of the underlying dynamical regime, in which trajectories transiently dwell near quasi-stable or saddle-like regions while retaining the capacity to move toward other configurations [41, 42]. Thus, spontaneous transitions between coordination states are observable consequences of metastability. However, such transitions alone do not define metastability [42]. In empirical EEG data, this underlying regime cannot be observed directly; instead, it is inferred from measurable properties, such as temporal variability in global synchronization and the diversity of spatial synchrony patterns. From this perspective, MSI and SCE provide complementary but partial views of metastable brain dynamics; MSI captures global synchronization variability, whereas SCE captures the repertoire of spatial phase synchronization patterns. This interpretation is consistent with the present results, in which SCE showed clearer sensor-frequency associations with AQ subscores, whereas MSI captured a weaker global variability component.

From a methodological perspective, the present results highlight important differences in the utility of MSI and SCE as metrics for metastable brain dynamics. Although both metrics are derived from phase synchronization, they capture complementary aspects of the network behavior. The MSI primarily reflects the temporal variability of global synchronization, providing a compact summary of how strongly brain regions fluctuate between more and less synchronized states. Therefore, MSI provides a simple summary of global coordination dynamics and offers a low-dimensional characterization of metastability. However, because MSI collapses spatial information into a single global measure, it may be less sensitive to differences in the configuration of network states over time. In contrast, SCE explicitly quantifies the diversity of the spatial patterns of phase synchronization, thereby capturing the repertoire of metastable network configurations. This may make SCE particularly sensitive to variations in the organization and reconfiguration of brain networks over time. The stronger and more consistent associations observed between SCE and AQ subscores in the current study suggest that individual differences in autistic traits are more closely related to the diversity of network states than to global synchronization variability. These distinctions indicate that SCE and MSI should not be considered interchangeable metrics of metastability but rather complementary metrics that emphasize different dimensions of brain dynamics. In particular, SCE may be more suitable for investigating cognitive and behavioral traits that depend on the flexible reconfiguration of large-scale networks, whereas MSI may be more appropriate for characterizing the overall stability–instability balance in neural synchronization. Therefore, the combined use of these metrics provides a more comprehensive framework for understanding metastable brain dynamics.

This study had several limitations. First, the present sample consisted of neurotypical adults and did not include participants clinically diagnosed with ASD. Therefore, the findings should be interpreted as trait-related associations rather than disease-specific neural markers. Second, the study was cross-sectional and correlational, and therefore cannot determine whether altered metastability-related dynamics causally contribute to autistic traits. Third, although cluster-based permutation tests controlled for multiple comparisons across electrodes and frequencies within each AQ measure, no additional correction was applied across the total AQ and all AQ subscores; these subscale-specific findings should therefore be interpreted as exploratory. Fourth, SCE depends on the phase-difference threshold used to define binary synchrony patterns.

Although the primary threshold was selected based on test–retest reliability, future studies should further evaluate the robustness of the findings across threshold choices and independent datasets. Fifth, the present analysis was conducted at the sensor level, and therefore the spatial interpretation of the topographic clusters should be treated cautiously. Finally, the UMAP analysis was based on illustrative individual examples and was not used for statistical inference. Future studies should quantify state-space properties across all participants and test whether these features generalize to independent resting-state and task-based EEG datasets.

In conclusion, our findings suggest that metastability metrics derived from EEG phase synchronization, particularly SCE, are associated with individual differences in autistic traits in a neurotypical cohort. These results highlight the relevance of spatiotemporal diversity in brain network dynamics in characterizing attention-switching– and communication-related autistic traits. While MSI showed additional, albeit weaker, associations, the combined use of complementary metrics offers a broader and more comprehensive characterization of metastable brain dynamics. This study contributes to a growing body of literature linking intrinsic neural dynamics to psychological traits within neurotypical populations and may provide a basis for future studies examining whether similar dynamical features are observed in clinically diagnosed ASD populations.

## Data and code availability

The resting-state EEG dataset analyzed in this study is publicly available on OSF: https://doi.org/10.17605/OSF.IO/29QB5. The MATLAB code used to compute the metastability index (MSI), channel-wise synchrony coalition entropy (SCE), and the statistical analyses is available at https://github.com/i-meb/EEG-Metastability. The repository includes example data, key parameter settings, and documentation for applying the analysis pipeline to preprocessed continuous EEG data.

## Supporting information

### Descriptive statistics

**S1 Table.**
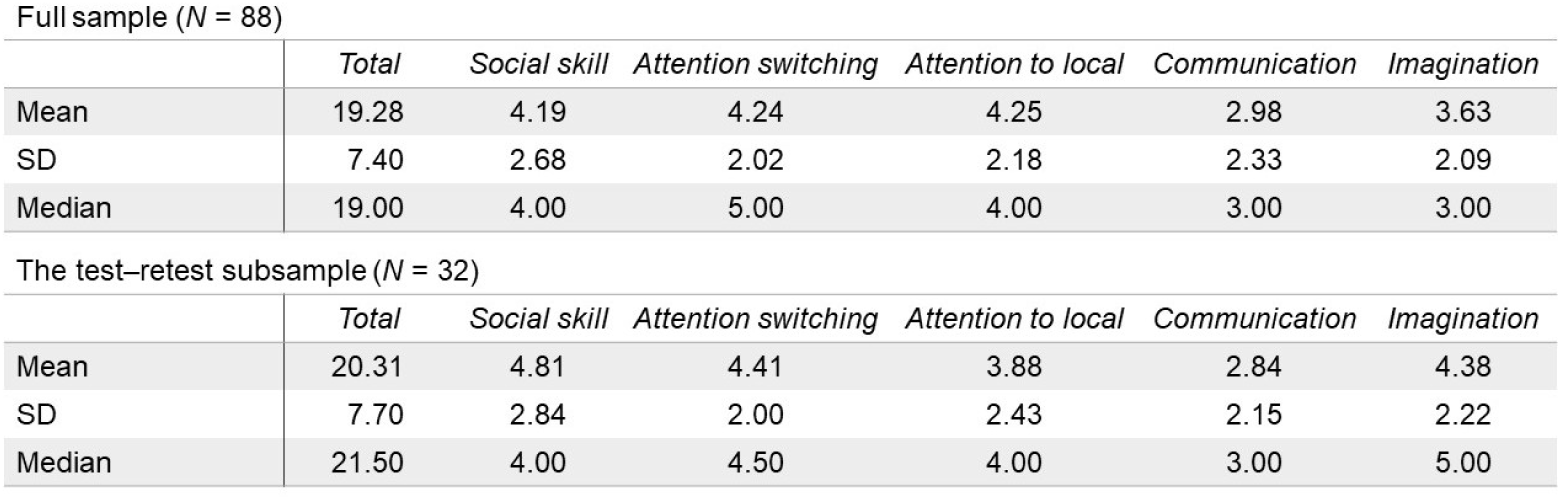
Descriptive statistics of the AQ scores of the 88 participants. S1 Table summarizes the descriptive statistics of AQ scores for both the full sample (*N* = 88) and the subsample that completed two days of sessions for the test–retest reliability assessment (*n* = 32). For each group, the mean, standard deviation (SD), median, and number of participants for the total AQ and each subscore (*social skill, attention switching, attention to detail, communication,* and *imagination*) were calculated.

**S1 Fig.**
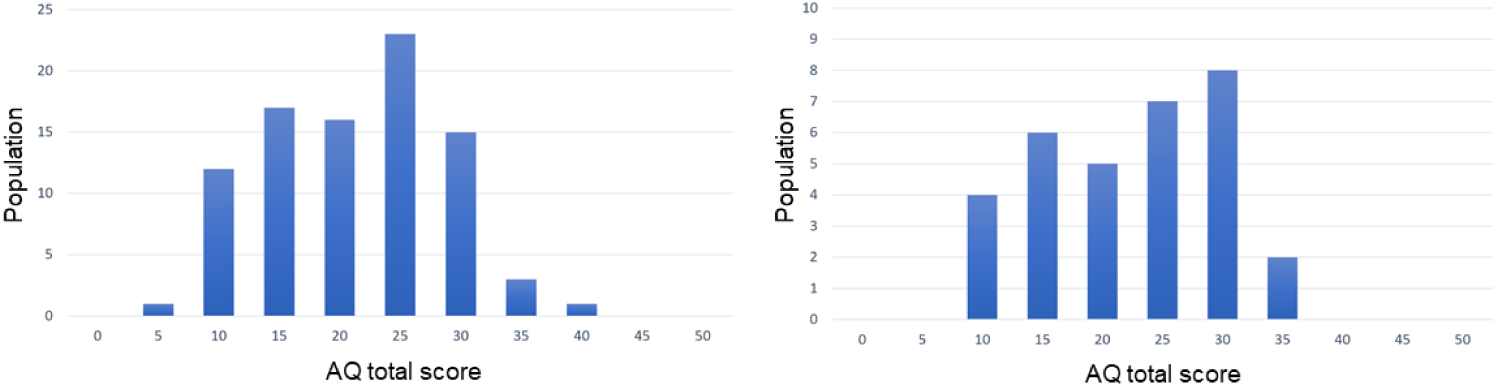
Distribution of the total AQ scores. For the full sample (*N* = 88; left panel of S1 Fig), the mean total AQ score was 19.28 (SD = 7.40), with a median of 19. The subscores ranged from 2.98 (*communication*) to 4.25 (*attention to detail*). In the test–retest subsample (*n* = 32, right panel of S1 Fig), the mean total AQ score was 20.31 (SD = 7.70), with a median of 21.5. The subscores ranged from 2.84 (*communication*) to 4.81 (*social skill*). The distribution of total AQ scores did not significantly deviate from normality in either the full sample (Shapiro–Wilk test, *p* = 0.206) or the test–retest subsample (*p* = 0.155).

### SCE thresholds determination

**S2 Fig.**
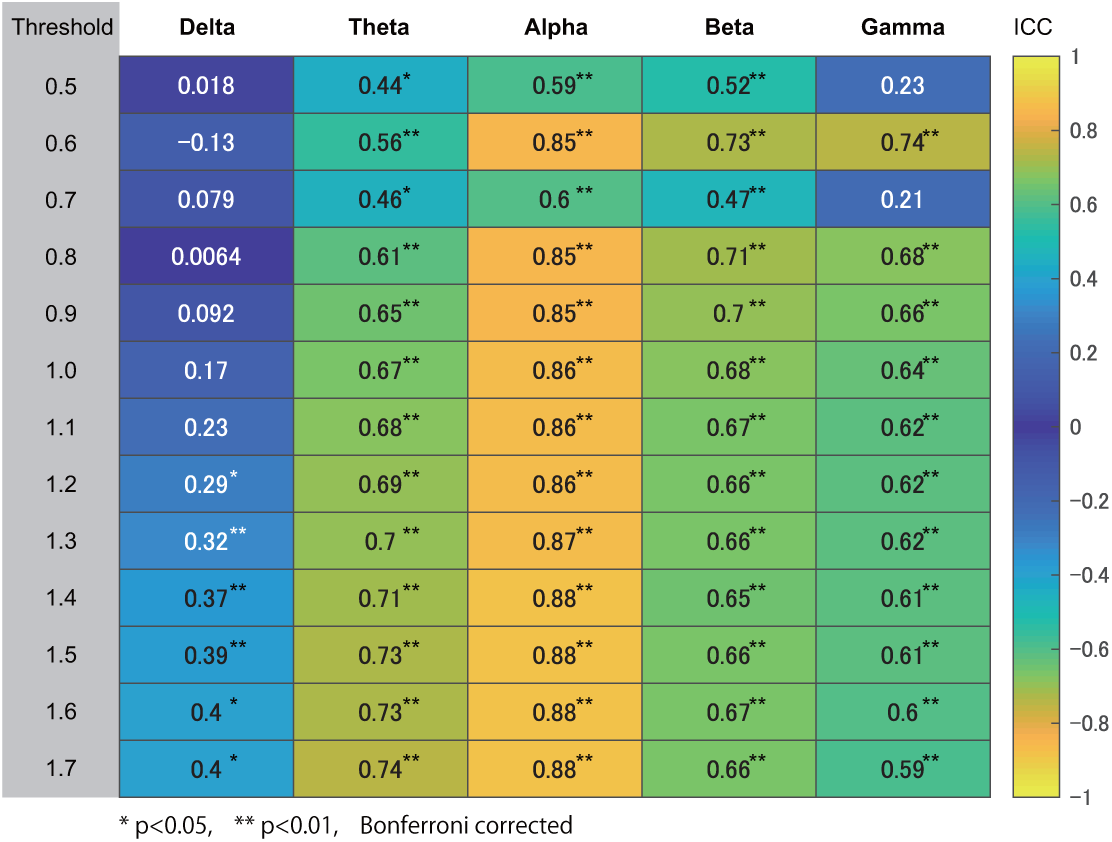
Threshold-dependent test–retest reliability of the SCE. A threshold of 0.8 rad has previously been proposed for binarizing instantaneous phase differences [15], but the validity of this threshold has not been systematically evaluated. Because the choice of threshold directly affects the binarization of phase differences and consequently the resulting SCE values, we systematically evaluated a range of threshold values. For each candidate threshold, SCE was calculated at the median frequency of each frequency band in the 32 participants who underwent follow-up measurements. We selected a threshold of 1.2 rad because it showed the highest overall test–retest reliability as assessed using the intraclass correlation coefficient (ICC). The threshold-dependent test–retest reliability of SCE across frequency bands is shown in S2 Fig. ICC values for each threshold (0.5–1.7 rad) and frequency band are shown in the heatmap. All p-values were Bonferroni-corrected for the number of frequency bands, with * indicating *p <* 0.05 and ** indicating *p <* 0.01.

### Correlations of MSI and oSCE with AQ

**S3 Fig.**
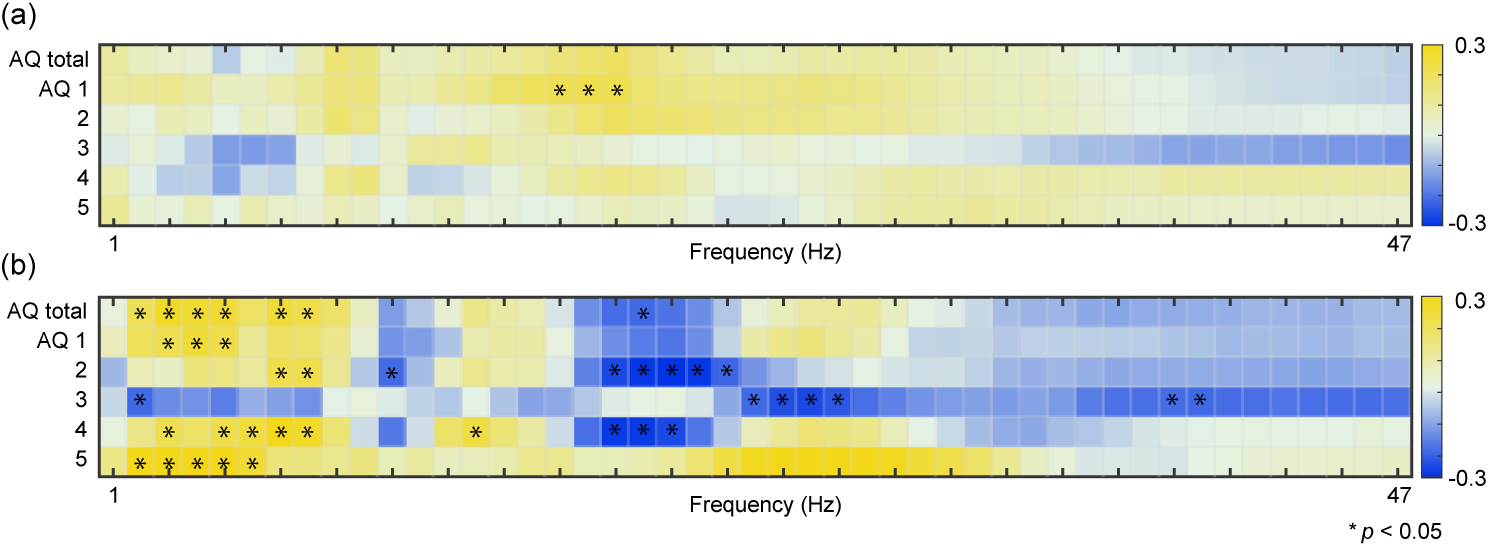
Frequency-wise correlations between AQ scores and whole-brain metastability metrics. Heatmaps show Spearman’s rank correlation coefficients between the total AQ score/AQ subscores and (a) MSI and (b) oSCE across 1–47 Hz. We found several positive correlations in the total AQ (2–5, 7–8, and 20 Hz), *social skill* (3–5 Hz), *attention switching* (7–8 Hz), *imagination* (3, 5–9, and 14 Hz), and *communication* (2–6 and 23–31 Hz). We also found some negative correlations in the total AQ (20 Hz), *attention switching* (11 and 19–23 Hz), *attention to detail* (2, 24–27, and 39–40 Hz), and *imagination* (19–21 Hz). Gray text indicates non-significant correlation coefficients. Bold text denotes statistically significant coefficients (*p <* 0.05) after the Bonferroni correction.

## References

1. Varela F, Lachaux JP, Rodriguez E, Martinerie J. The brainweb: Phase synchronization and large-scale integration. Nature Reviews Neuroscience. 2001;2(4):229–39. doi:10.1038/35067550.

2. Tognoli E, Kelso JAS. The metastable brain. Neuron. 2014;81(1):35-48. doi:10.1016/j.neuron.2013.12.022.

3. Kelso JAS. Dynamic patterns: the self-organization of brain and behavior. Cambridge, MA: MIT Press; 1995.

4. Kelso JAS. Multistability and metastability: Understanding dynamic coordination in the brain. Philosophical Transactions of the Royal Society B: Biological Sciences. 2012;367(1591):906–18. doi:10.1098/rstb.2011.0351.

5. Deco G, Tononi G, Boly M, Kringelbach ML. Rethinking segregation and integration: Contributions of whole-brain modelling. Nature Reviews Neuroscience. 2015;16(7):430–9. doi:10.1038/nrn3963.

6. Deco G, Kringelbach ML, Jirsa VK, Ritter P. The dynamics of resting fluctuations in the brain: Metastability and its dynamical cortical core. Scientific Reports. 2017;7(1):3095. doi:10.1038/s41598-017-03073-5.

7. Brock ME, Freuler A, Baranek GT, Watson LR, Poe MD, Sabatino A. Temperament and sensory features of children with autism. Journal of Autism and Developmental Disorders. 2012;42(11):2271–84. doi:10.1007/s10803-012-1472-5.

8. Robinson EB, Munir K, Munafò MR, Hughes M, McCormick MC, Koenen KC. Stability of autistic traits in the general population: Further evidence for a continuum of impairment. Journal of the American Academy of Child & Adolescent Psychiatry. 2011;50(4):376–84. doi:10.1016/j.jaac.2011.01.005.

9. Palmer CJ, Paton B, Enticott PG, Hohwy J. ’Subtypes’ in the presentation of autistic traits in the general adult population. Journal of Autism and Developmental Disorders. 2015;45(5):1291–301. doi:10.1007/s10803-014-2289-1.

10. Constantino JN, Todd RD. Autistic traits in the general population: A twin study. Archives of General Psychiatry. 2003;60(5):524. doi:10.1001/archpsyc.60.5.524.

11. Michel CM, Koenig T. EEG microstates as a tool for studying the temporal dynamics of whole-brain neuronal networks: A review. NeuroImage. 2018;180:577–93. doi:10.1016/j.neuroimage.2017.11.062.

12. Seeber M, Michel CM. Synchronous brain dynamics establish brief states of communality in distant neuronal populations. eNeuro. 2021;8(3):ENEURO.0005-21.2021. doi:10.1523/ENEURO.0005-21.2021.

13. Sase T, Kitajo K. The metastable brain associated with autistic-like traits of typically developing individuals. PLOS Computational Biology. 2021;17(4):e1008929. doi:10.1371/journal.pcbi.1008929.

14. Jia H, Gao F, Yu D. Altered temporal structure of neural phase synchrony in patients with autism spectrum disorder. Frontiers in Psychiatry. 2021;12:618573. doi:10.3389/fpsyt.2021.618573.

15. Shanahan M. Metastable chimera states in community-structured oscillator networks. Chaos: An Interdisciplinary Journal of Nonlinear Science. 2010;20(1):013108. doi:10.1063/1.3305451.

16. Schartner M, Seth A, Noirhomme Q, Boly M, Bruno MA, Laureys S, et al. Complexity of multi-dimensional spontaneous EEG decreases during propofol induced general anaesthesia. PLOS ONE. 2015;10(8):e0133532. doi:10.1371/journal.pone.0133532.

17. Baron-Cohen S, Wheelwright S, Skinner R, Martin J, Clubley E. The autism-spectrum quotient (AQ): Evidence from Asperger syndrome/high-functioning autism, males and females, scientists and mathematicians. Journal of Autism and Developmental Disorders. 2001;31(1):5–17. doi:10.1023/A:1005653411471.

18. Wakabayashi A, Baron-Cohen S, Wheelwright S, Tojo Y. The autism-spectrum quotient (AQ) in Japan: A cross-cultural comparison. Journal of Autism and Developmental Disorders. 2006;36(2):263–70. doi:10.1007/s10803-005-0061-2.

19. Wakabayashi A, Baron-Cohen S, Uchiyama T, Yoshida Y, Tojo Y, Kuroda M, et al. The autism-spectrum quotient (AQ) children’s version in Japan: A cross-cultural comparison. Journal of Autism and Developmental Disorders. 2007;37(3):491–500. doi:10.1007/s10803-006-0181-3.

20. Delorme A, Makeig S. EEGLAB: An open source toolbox for analysis of single-trial EEG dynamics including independent component analysis. Journal of Neuroscience Methods. 2004;134(1):9–21. doi:10.1016/j.jneumeth.2003.10.009.

21. Oostenveld R, Fries P, Maris E, Schoffelen JM. FieldTrip: Open source software for advanced analysis of MEG, EEG, and invasive electrophysiological data. Computational Intelligence and Neuroscience. 2011;2011:1–9. doi:10.1155/2011/156869.

22. Delorme A, Sejnowski T, Makeig S. Enhanced detection of artifacts in EEG data using higher-order statistics and independent component analysis. NeuroImage. 2007;34(4):1443–9. doi:10.1016/j.neuroimage.2006.11.004.

23. Pion-Tonachini L, Kreutz-Delgado K, Makeig S. ICLabel: An automated electroencephalographic independent component classifier, dataset, and website. NeuroImage. 2019;198:181–97. doi:10.1016/j.neuroimage.2019.05.026.

24. Kayser J, Tenke CE. Principal components analysis of Laplacian waveforms as a generic method for identifying ERP generator patterns: I. Evaluation with auditory oddball tasks. Clinical Neurophysiology. 2006;117(2):348–68. doi:10.1016/j.clinph.2005.08.034.

25. Kayser J, Tenke CE. Principal components analysis of Laplacian waveforms as a generic method for identifying ERP generator patterns: II. Adequacy of low-density estimates. Clinical Neurophysiology. 2006;117(2):369–80. doi:10.1016/j.clinph.2005.08.033.

26. Lachaux JP, Rodriguez E, Martinerie J, Varela FJ. Measuring phase synchrony in brain signals. Human Brain Mapping. 1999;8(4):194–208. doi:10.1002/(SICI)1097-0193(1999)8:4¡194::AID-HBM4¿3.0.CO;2-C.

27. Cohen MX. Analyzing neural time series data: Theory and practice. Cambridge, MA: MIT Press; 2014.

28. Kuramoto Y. Chemical oscillations, waves, and turbulence. vol. 19 of Springer Series in Synergetics. Berlin, Heidelberg: Springer Berlin Heidelberg; 1984. doi:10.1007/978-3-642-69689-3.

29. McInnes L, Healy J, Melville J. UMAP: uniform manifold approximation and projection for dimension reduction. arXiv preprint arXiv:180203426. 2018. doi:10.48550/arXiv.1802.03426.

30. Meehan C, Ebrahimian J, Moore W, Meehan S. Uniform manifold approximation and projection (UMAP); 2025. MATLAB Central File Exchange. Available from: https://www.mathworks.com/matlabcentral/fileexchange/71902.

31. Geurts HM, Corbett B, Solomon M. The paradox of cognitive flexibility in autism. Trends in Cognitive Sciences. 2009;13(2):74–82. doi:10.1016/j.tics.2008.11.006.

32. Miller HL, Ragozzino ME, Cook EH, Sweeney JA, Mosconi MW. Cognitive set shifting deficits and their relationship to repetitive behaviors in autism spectrum disorder. Journal of Autism and Developmental Disorders. 2015;45(3):805–15. doi:10.1007/s10803-014-2244-1.

33. Lage C, Smith ES, Lawson RP. A meta-analysis of cognitive flexibility in autism spectrum disorder. Neuroscience & Biobehavioral Reviews. 2024;157:105511. doi:10.1016/j.neubiorev.2023.105511.

34. Engel AK, Fries P. Beta-band oscillations — signalling the status quo? Current Opinion in Neurobiology. 2010;20(2):156–65. doi:10.1016/j.conb.2010.02.015.

35. Geurts HM, Verté S, Oosterlaan J, Roeyers H, Sergeant JA. How specific are executive functioning deficits in attention deficit hyperactivity disorder and autism? Journal of Child Psychology and Psychiatry. 2004;45(4):836–54. doi:10.1111/j.1469-7610.2004.00276.x.

36. Dajani DR, Uddin LQ. Demystifying cognitive flexibility: Implications for clinical and developmental neuroscience. Trends in Neurosciences. 2015;38(9):571–8. doi:10.1016/j.tins.2015.07.003.

37. Bastiaansen M, Hagoort P. Oscillatory neuronal dynamics during language comprehension. In: Progress in Brain Research. vol. 159. Elsevier; 2006. p. 179–96. doi:10.1016/S0079-6123(06)59012-0.

38. Jochaut D, Lehongre K, Saitovitch A, Devauchelle AD, Olasagasti I, Chabane N, et al. Atypical coordination of cortical oscillations in response to speech in autism. Frontiers in Human Neuroscience. 2015;9. doi:10.3389/fnhum.2015.00171.

39. Pavlova MA. Biological motion processing as a hallmark of social cognition. Cerebral Cortex. 2012;22(5):981–95. doi:10.1093/cercor/bhr156.

40. Oberman LM, Hubbard EM, McCleery JP, Altschuler EL, Ramachandran VS, Pineda JA. EEG evidence for mirror neuron dysfunction in autism spectrum disorders. Cognitive Brain Research. 2005;24(2):190–8. doi:10.1016/j.cogbrainres.2005.01.014.

41. Heitmann S, Breakspear M. Putting the “dynamic” back into dynamic functional connectivity. Network Neuroscience. 2018 Jan;2(2):150–74. doi:10.1162/netn_a_00041.

42. Hancock F, Rosas FE, Luppi AI, Zhang M, Mediano PAM, Cabral J, et al. Metastability demystified — The foundational past, the pragmatic present and the promising future. Nature Reviews Neuroscience. 2025;26(2):82–100. doi:10.1038/s41583-024-00883-1.

